# Consequences of cannibalism: induced defense and kin discrimination in a rotifer

**DOI:** 10.1101/2025.10.31.685857

**Authors:** Emily A. Harmon, Alisher K. Bimagambetov, Henry T. Lee, David W. Pfennig

## Abstract

Resource polyphenisms result in the production of environmentally induced morphs with differential niche use. These striking examples of phenotypic plasticity are taxonomically widespread and ecologically important. However, cannibalism is a frequent repercussion of resource polyphenisms. Despite some benefits, cannibalism is always costly to victims and sometimes costly to the cannibal. Therefore, we here evaluate how morphological and behavioral plastic strategies to minimize these costs may evolve. To better understand the evolution and consequences of cannibalistic polyphenisms, we tested whether rotifers *Asplanchna brightwellii* possess effective morphological defenses against cannibalism and the ability to discriminate by genetic relationship. We found that small humps produced by vulnerable *A. brightwellii* limit cannibalism. We also found that cannibals were less likely to attack clonemates than non-kin. The rotifer genus *Asplanchna* comprises species with varying degrees of resource polyphenism, cannibalism, induced morphological defenses, and behavioral kin discrimination. The observed induced defense and kin discrimination in *Asplanchna brightwellii* likely represent evolutionary intermediates facilitating the evolution of a unique trimorphic resource polyphenism in congeners.

## Introduction

Diverse taxa can alter their behavior, physiology, and morphology in response to environmental variation via phenotypic plasticity. A common and often ecologically important form of plasticity is resource polyphenism, in which distinct morphs specializing on alternate resources co-occur in a population. A resource polyphenism can allow organisms to match currently available resources or partition resources within and between species [1, 2]. Morphs that feed at a higher trophic level tend to have larger feeding apparatus and body size, thereby consuming larger food including intraguild competitors [3]. However, by enlarging gape-size it becomes possible to consume not only larger prey, but also conspecifics [4, 5]. Consequently, cannibalistic polyphenisms are widespread across taxa from protozoans to amphibians [e.g., 3, 6, 7-10].

Cannibalism has various consequences. The cannibal gains a benefit by simultaneously receiving a meal and eliminating a competitor [9, 11]. However, there is a high cost to the victim: death. One potential consequence of cannibalism is therefore the evolution of induced defenses, such as shortening vulnerable life periods, avoiding potential cannibals, or changing morphology in environments where cannibalism is likely [3, 12-14]. Cannibalism can also be costly to the cannibals: if the victim is a close relative, the cannibal incurs inclusive fitness costs [15]. Inclusive fitness encompasses the ability of an organism to increase its genetic fitness either directly by transmitting alleles through offspring or indirectly by enhancing the survival of its relatives, which carry a portion of its alleles [16]. Sophisticated behaviors such as kin discrimination can evolve to limit inclusive fitness costs, allowing cannibals to differentially interact with individuals based on genetic relatedness [17-22].

Here, we evaluate the morphological and behavioral consequences of cannibalism in an organism with a cannibalistic resource polyphenism. We predict that under environmental conditions in which the threat of cannibalism is high (i.e., environments that induce the cannibalistic morph), plastic morphological and/or behavioral responses to cannibalism will be expressed. Indeed, plastic responses to cannibalism may be especially likely to evolve in cannibalistic polyphenisms, which already exhibit plastic variation in morphology and behavior.

We evaluate the consequences of cannibalism in *Asplanchna*, a genus of predatory rotifers. *Asplanchna* species generally fall into two groups: small, monomorphic species that eat phytoplankton and small rotifers [23-27], or species with a highly developed trimorphic resource polyphenism (Figure 1). The morphs in the trimorphic species are 1) a small sac-shaped (‘saccate’) morph, similar to monomorphic *Asplanchna*, 2) an intermediate-sized, humped (‘cruciform’) morph, and 3) a large, bell-shaped (‘campanulate’), particularly predatory morph [28-30]. The two larger morphs can eat larger prey and specifically prefer and perform best on a diet of congeneric or conspecific prey [31, 32]. In fact, in a field experiment, the majority of sampled individuals of the trimorphic species *Asplanchna sieboldii* had conspecific remains in their stomachs [33].

**Figure 1.**
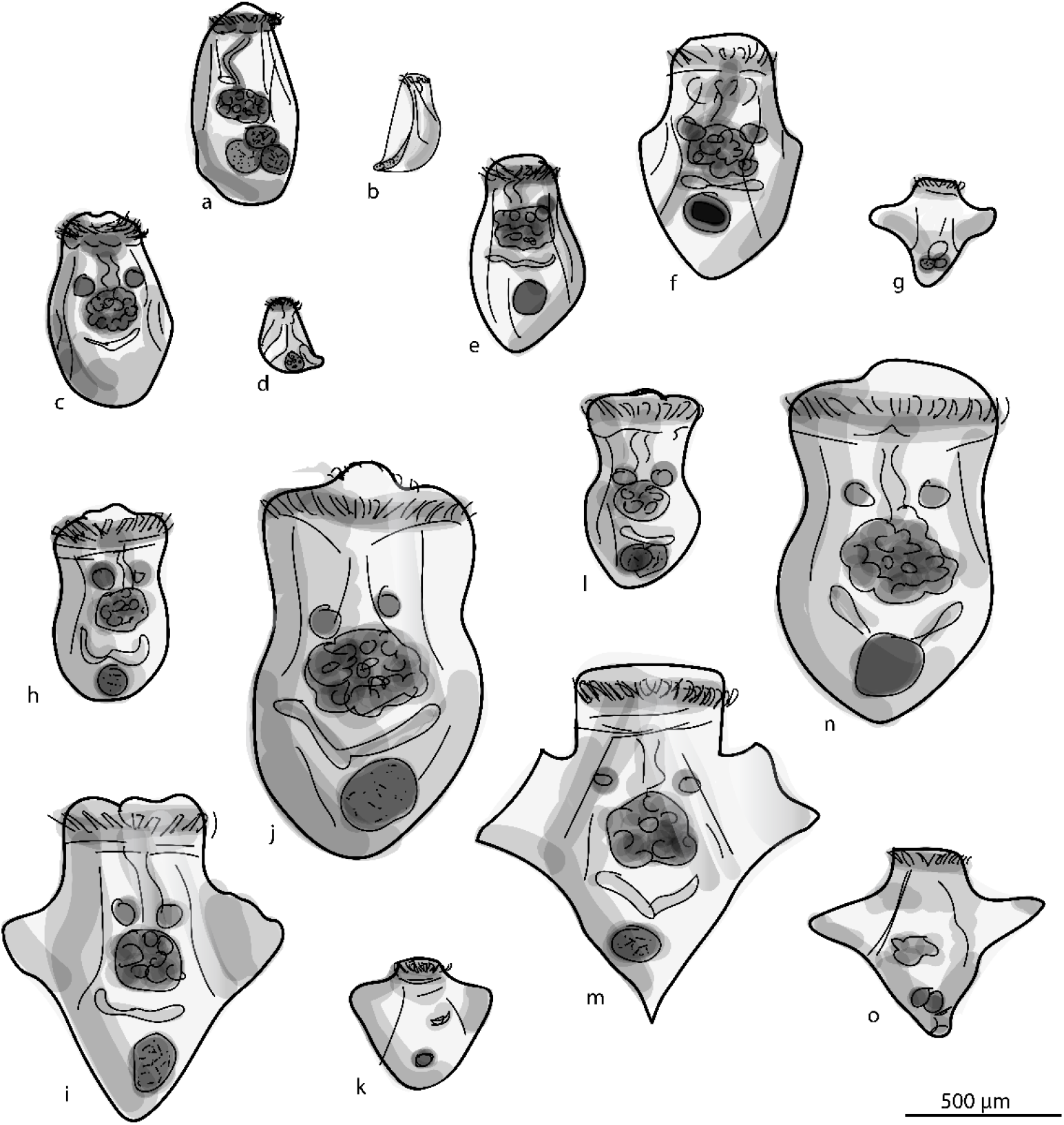
A selection of morphology across *Asplanchna*. Species with monomorphic females: (a) *A. priodonta* female and (b) male, (c) *A. girodi* female and (d) male. Weak polyphenism in *A. brightwellii*: (e) alpha morph, (f) beta morph, (g) male. Species with trimorphic females: (h) *A. intermedia* primary saccate morph, (i) intermediate cruciform morph, (j) campanulate morph, and (k) male. (l) *A. sieboldii* primary saccate morph, (m) intermediate cruciform morph, (n) campanulate morph, and (o) male. Drawings are to scale with average mature body sizes.

To better understand the consequences of cannibalistic polyphenisms, we focus on *Asplanchna brightwellii*, which exhibit an intermediate polyphenism (Figure 1) and may represent early evolutionary stages of the complex trimorphic polyphenism [34]. *A. brightwellii* do not produce the campanulate morph, just a small, sac-shaped morph (‘alpha’) and a larger, humped morph (‘beta’). The size difference between the morphs is more subtle than in the trimorphic species – the beta morph is on average 1.5x larger than the alpha morph, relative to the 1.6-2x size difference across *A. sieboldii* morphs [35-37]. The female humps are also smaller than in the trimorphic species – beta morphs have a dorsal hump and sometimes small lateral humps [38]. *A. brightwellii* consume prey of an intermediate size and are only slightly cannibalistic; in nature, 3% or less of *A. brightwellii* were found with conspecifics in their stomachs [25, 39-42].

We first examine induced defenses. Following our above prediction, we expect to see a morphological defense in *Asplanchna* species that have a cannibal morph, and that the defense is expressed under conditions that induce the cannibal morph. The humps present in some *Asplanchna* are likely a defense against cannibalism. Humped individuals have up to four body-wall outgrowths that are extended with hydrostatic pressure, increasing their body size so that they are more difficult to eat [25, 33, 43-45]. In a direct test, J.J. Gilbert showed that of 93 options of each prey type, 43 non-humped females (46%) and only 7 humped females (8%) were consumed by *A. sieboldii* cannibals [25]. Critically, only males and females of the polyphenic species have humps, and these humps are only expressed under environmental conditions in which the most cannibalistic individuals are induced (Table 1). Further, across the genus, the more cannibalistic species tend to have more pronounced humps [Table 1; Figure 1; 25, 34]. However, it is unknown whether the humps of the more weakly polyphenic species, *A. brightwellii*, deter cannibalism.

**Table 1.** Literature survey of cannibalism, polyphenism, defenses, and potential kin discrimination across the genus *Asplanchna*. Species are listed in increasing order of tendency to take large metazoan prey [42].

| Species | Predominate food | Cannibalistic | Resource polyphenism | Male humps | Kin discrimination | Sources |
| --- | --- | --- | --- | --- | --- | --- |
| *A. priodonta* | Phytoplankton | No | No | No | - | [25, 41, 57-65] |
| *A. herrickii* | Phytoplankton<br>Small rotifers | No | No | No | - | [25, 57, 59, 66] |
| *A. tropica* | Small rotifers* | No | No | ? | - | [27, 67] |
| *A. asymmetrica* | Small rotifers* | No | No | ? | - | [68] |
| *A. girodi* | Phytoplankton<br>Small rotifers | No | No | No | - | [25, 41, 57, 60, 64, 69, 70] |
| *A. brightwellii* | Rotifers | Yes | Dimorphic (small humps) | Moderate | ? | [25, 39-41, 71, 72] |
| *A. intermedia* | Rotifers* | Yes | Trimorphic (moderate humps) | Moderate | Highly selective species discrimination | [44, 45, 73, 74] |
| *A. sieboldii* | Large rotifers<br>Crustaceans | Yes | Trimorphic (large humps) | Large | Somewhat selective species discrimination | [25, 33, 41, 44, 45] |
| *A. silvestrii* | Large rotifers* | Yes | Trimorphic (large humps) | Yes | ? | [25, 29, 41, 75] |

\*Few studies of stomach contents in nature

We next evaluate evidence for behavioral plasticity in response to cannibalism: kin discrimination. It has been demonstrated that *A. intermedia* and *A. sieboldii* consume more congenerics than clonemates [43-45]. These cannibals can also discriminate between different morphs and sexes within their own lineage, preferentially eating saccate females [44, 45]. However, given that just one clonal lineage of each species has been tested, it is not clear whether the mechanism of discrimination was species-recognition, kin-recognition, or self-recognition. It is also unknown whether other species of *Asplanchna* exhibit any form of kin discrimination.

Here we test whether the slight humps present in beta morphs of *A. brightwellii* are protective against cannibalism and whether there is evidence for kin discrimination. We evaluate multiple clonal lineages to obtain patterns that are more generalizable across the species. By evaluating these consequences of cannibalism in a species representing the early evolutionary stages of a complex cannibalistic polyphenism, we can better understand the evolution of resource polyphenism, cannibalism, induced defenses, and kin discrimination.

## Methods and Materials

Stock cultures of *A. brightwellii* from five Texas ponds were obtained and cultured as described elsewhere [37, 46]. A clonal culture from Greenville, North Carolina was started from a single female isolated from a pond. We induced bisexual stocks of beta morphs by periodically adding D*-*α-tocopherol (vitamin E; the dietary cue that induces the beta morphotype and sexual reproduction) to an approximate concentration of 10^-7^ M. Stock cultures were fed *Paramecium aurelia* during the test of hump function and were fed *Philodina* spp. (which was found to support more stable cultures) during the kin recognition tests. *Philodina* spp. were cultured on nutritional yeast.

### Induced defense

We performed prey choice tests to evaluate whether humps on beta females are protective against cannibalism in *A. brightwellii*. We placed mature beta morphs (predators) with equal numbers of small size-matched alpha and beta females (prey) to evaluate if females with humps (beta morphs) were less likely to be consumed than those without humps (alpha morphs).

We obtained neonates by isolating 20 adults from a stock culture in a Petri dish filled with EPA medium [47]. After four to six hours, we removed neonates that had been produced. To size-match rotifers, we photographed them using the 4x objective of an Olympus CX43 microscope with the Olympus LC30 camera and 0.5x adapter. We measured body lengths in Olympus cellSens imaging software. Due to beta neonates being born at a relatively large size (Table S1), we size-matched alpha adults with beta neonates. Across the 13 trials, beta predators were sourced from four different ponds, beta prey were sourced from three ponds, and alpha prey were sourced from four clonal lineages. In each trial, predator and prey were sourced from different ponds to control for possible effects of kin recognition.

We conducted prey choice tests in the wells of a 12-well culture plate filled halfway with EPA medium. Each well contained 3-8 predators from paramecia-depleted stock cultures and equal numbers (3-8) of each prey type. After one hour, we fixed all rotifers in ethanol to expand their humps and counted the remaining number of each prey type. One hour was chosen to minimize the birth of additional neonates.

All analyses were conducted in R version 4.3.3 [48]. We ran a generalized linear mixed effects model with a binomial distribution to assess whether the probability of being eaten varied between humped and non-humped females [glmer in ‘lme4’; 49]. Trial was included as a random intercept. We tested the effect of morph with a likelihood ratio test and checked model assumptions using package ‘DHARMa’ [50].

### Kin discrimination

We performed prey choice tests to evaluate whether clonemates were less likely to be cannibalized. We placed mature beta females (predators) with equal numbers of small size-matched kin (clonemates) and non-kin alpha morphs. We distinguished lineages by staining either the kin or non-kin lineage with 0.5 µg/mL of neutral red chloride. This treatment was previously found to have no adverse effect on *A. brightwellii* [51]. The staining of *A. brightwellii* was visible to the naked eye within 12 hours and lasted for approximately three days without additional treatment.

We tested prey selection in three different clonal lineages, each from a different pond: Laguna Prieta (GPS coordinates 31.9247, -106.0471), Behind Ranch House (31.9241, - 106.0417), and Brody Duck Pond (35.6125, -77.4029). Replicate trials for each predator were performed alternating whether the kin or the non-kin prey were stained. Each predator was tested against two potential non-kin lineages. Each combination of participants was replicated three times for a total of 36 trials (*N* = 3 predator lineages x 2 non-kin lineages x 2 stain combinations x 3 replicates = 36).

We conducted choice tests in the wells of a 24-well culture plate containing about 0.2 mL of EPA medium. In each trial, we placed a mature beta female from a *Philodina*-depleted stock culture with five alpha clonemates and five alpha non-kin clones from a different pond. For 10 minutes, we observed the feeding behavior of the beta morph under a stereomicroscope. We counted encounters, attacks, captures, and ingestions of prey. An encounter was classified as a “bump”, or passing by with contact between the two morphs while an attack occurred when the corona of the beta morph widened in an attempt to engulf the alpha morph. A capture was recorded when the beta successfully engulfed the alpha. A capture without release was an ingestion. After the 10 minutes of observation, trials were run for an additional 30 minutes and the remaining prey were counted.

We first analyzed whether the probability of being cannibalized depended upon kin-status. We ran a generalized linear mixed effects model with a binomial distribution and fixed effect of kin-status. Participant combination (prey and predator lineages, kin or non-kin stained) was included as a random intercept. We next ran similar models using the response variables of attack probability (the proportion of encounters leading to an attack) and capture probability (the proportion of attacks leading to a capture) to evaluate if cannibals differed in feeding behavior toward the different kin types. We used likelihood ratio tests to evaluate the fixed effects and checked model assumptions using package ‘DHARMa’ [50].

## Results

### Induced defense

Beta neonates (females with humps) were less likely to be cannibalized than alpha adults (females without humps) of the same size (Likelihood ratio test [LRT] = 4.18, df = 1, *p* = 0.041). Of the 72 options of each prey type, 18 alpha morphs and 9 beta morphs were consumed across 13 trials (Figure 2a).

**Figure 2.**
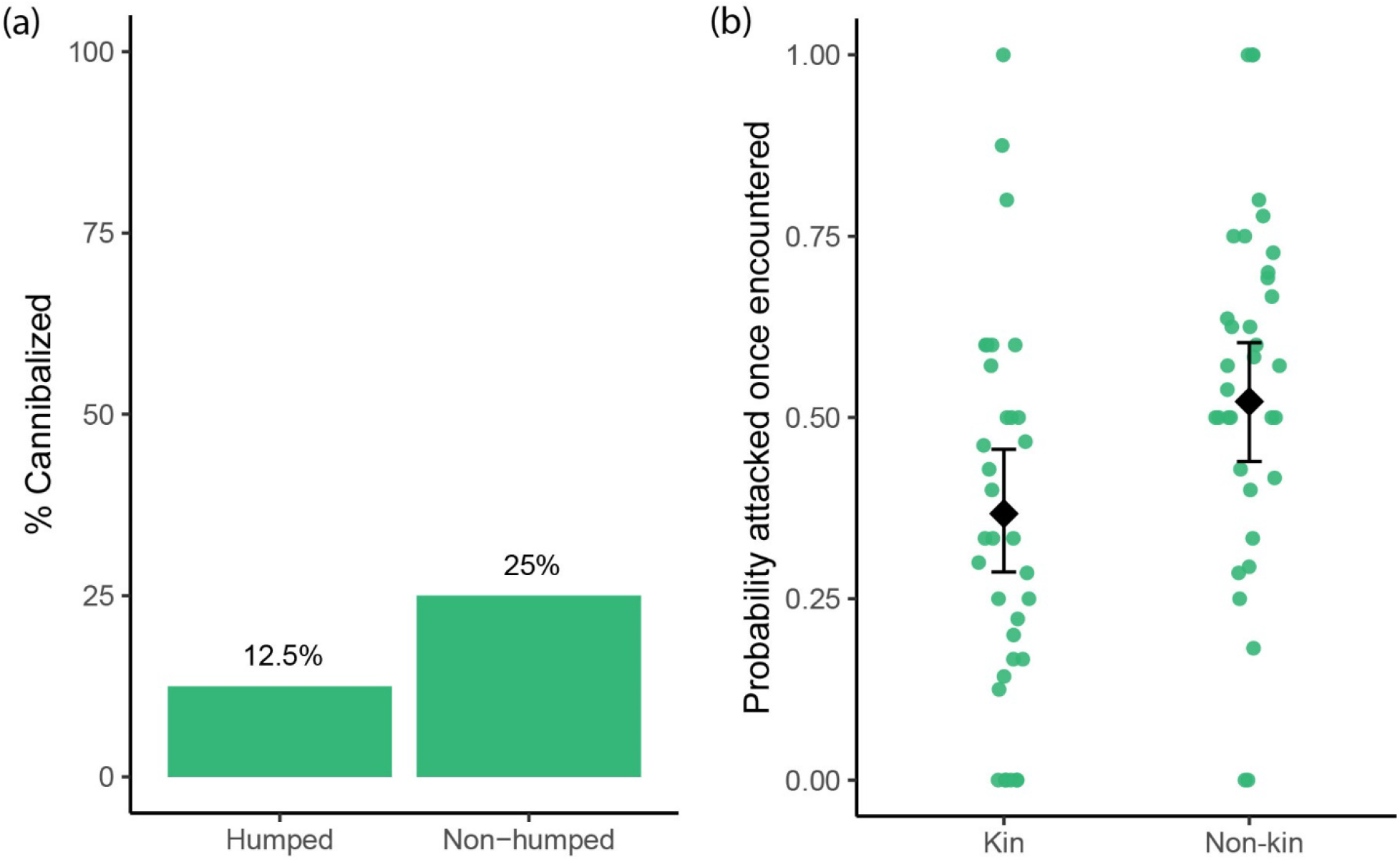
*A. brightwellii* defenses against cannibalism. (a) Percentage of size-matched humped and non-humped *A. brightwellii* that were eaten by mature beta morphs across 13 trials. (b) Mature beta morphs given a choice of prey between kin (clonemates) and non-kin (lineage from another pond) alpha morphs. Points show the probability of attack for each trial. Black diamond and bars show estimate and 95% confidence intervals from logistic regression.

### Kin discrimination

Only 14 of the 360 potential prey were consumed (5 kin, 10 non-kin; 6 control, 9 dyed). Kin status (LRT = 1.79, df = 1, *p* = 0.18) did not impact the likelihood of being eaten. However, feeding behavior of potential cannibals varied with prey relationship (Table 2). Upon encountering a rotifer, beta morphs were more likely to attack non-kin rotifers than they were to attack clonemates (Figure 2b; LRT = 10.7, df = 1, *p* = 0.0011). Once an alpha morph was attacked, the likelihood that it was captured did not vary according to relationship (LRT = 0.23, df = 1, *p* = 0.63). Only three alpha rotifers were ingested during the behavioral observations.

**Table 2.**
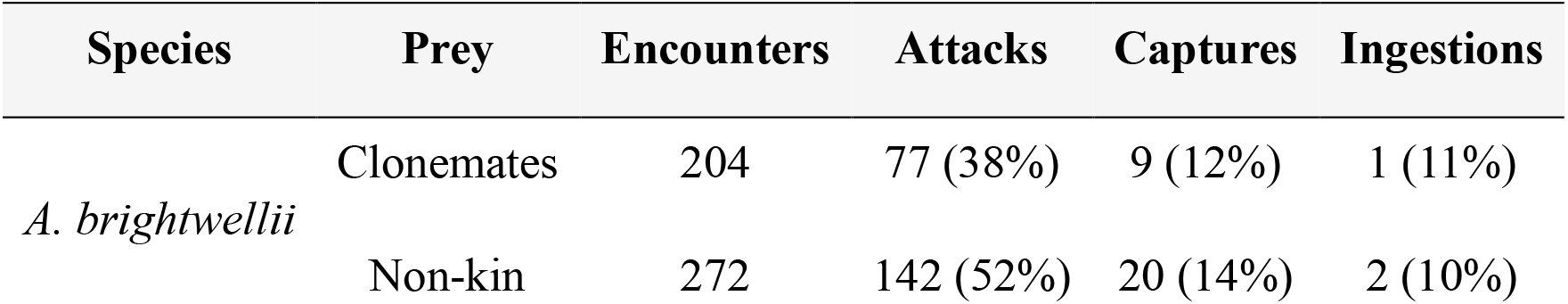
Feeding behavior of *A. brightwellii* beta morphs presented with size-matched alpha morphs of the same or different lineage. Summary of 36 trials.

| Species | Prey | Encounters | Attacks | Captures | Ingestions |
| --- | --- | --- | --- | --- | --- |
| *A. brightwellii* | Clonemates | 204 | 77 (38%) | 9 (12%) | 1 (11%) |
|  | Non-kin | 272 | 142 (52%) | 20 (14%) | 2 (10%) |

## Discussion

Understanding the consequences of cannibalism in *A. brightwellii* informs hypotheses for the evolution of resource polyphenism in *Asplanchna* [34]. The ancestral *Asplanchna* was likely monomorphic. Across Rotifera, body size is a variable and plastic trait influenced by factors such as temperature, prey size, maternal size, and genetics [35, 52-54]. The largest individuals of ancestral *Asplanchna* could take advantage of large prey items when they are available. Consuming a higher trophic level (including competitors) would be favored under conditions in which there is high competition for small prey. As the diet contained more large, herbivorous prey than small prey such as ciliates and detritivores, vitamin E from phytoplankton would accumulate in body tissues. *Asplanchna* evolved to respond to high dietary vitamin E (and the presence of large prey) by producing larger, more carnivorous offspring [28, 34, 36].

Following the evolution of the growth response to vitamin E, cannibalism and congeneric predation would have become an issue. Lineages in which vitamin E induced cell growth disproportionally in some regions of the hypodermis [55] would become a shape harder to consume by conspecifics. This would lead to the evolution of humps which protected young and males from cannibalism and congeneric predation [34]. Our results in *A. brightwellii* are consistent with these hypotheses: alpha adults were twice as likely to be cannibalized as size-matched beta neonates. Alpha and beta morphs likely do not coexist in nature for long; when threshold vitamin E levels are reached, lineages fully transition from producing alpha morphs to beta morphs within a few generations [35]. Humps on beta morphs can therefore effectively minimize cannibalism.

Additionally, differences would have arisen between lineages in the likelihood of being a cannibal depending on factors such as size, population dynamics, or prey availability. Even if humps had already evolved, some lineages would have a higher chance of eating relatives than others. Kin selection would favor the evolution of kin discrimination in lineages with a high risk of cannibalism. We found that there is kin discrimination in *A. brightwellii;* beta morphs can distinguish clonemates from others of their species, preferentially attacking non-clones (Table 2). Further work can investigate whether this discrimination is a form of self-recognition (clones vs. non-clones) or scales with degree of genetic distance within and across species.

Once induced defenses and kin discrimination such as in *A. brightwellii* had evolved, it would be easier for the full trimorphic expression of the resource polyphenism to evolve. Neither strategy to avoid cannibalism appears to be particularly costly– humps are not extended while swimming so they have little hydrodynamic effect [29], and kin discrimination can be overruled when it is less beneficial [e.g., 56]. On the other hand, cannibalism can impose high costs to victims and cannibals; even with evolved defenses in place, high rates of cannibalism can occur [33]. Especially as the growth response evolved to become more extreme, cannibalism would be a strong selective force in *Asplanchna*. Adaptive refinement of humps and kin discrimination could have led to the extreme polyphenisms seen in *A. intermedia, A. sieboldii*, and *A. silvestrii*.

Finally, variation in the risk of cannibalization across lineages could lead to different patterns of selection on humps versus kin discrimination. These two consequences of cannibalism may evolve reciprocally: a lineage with very pronounced humps may not evolve particularly selective kin discrimination and vice versa. Moreover, selection on induced defenses and kin discrimination likely varies across environments – especially those with varying threats from congeneric versus conspecific predation. For instance, in sympatric populations, humps would protect from both conspecific and congeneric predation and may be the predominate solution to cannibalism. Alternatively, kin discrimination may be the more dominant strategy if it has additional benefits such as facilitating outcrossing during bisexual reproduction. Evidence for a tradeoff is still limited: *A. sieboldii* are highly defended and moderately discriminative, *A. intermedia* are moderately defended and highly discriminative, while *A. brightwellii* are slightly defended and slightly discriminative. More lineages will need to be evaluated to clarify if there is a relationship between kin discrimination and induced defenses.

Further investigations of traits across *Asplanchna* will help us continue to parse out how the trimorphic polyphenism evolved. *Asplanchna* remain an underexplored but promising system in which to evaluate not just the evolution of polyphenisms, but also cannibalism and kin recognition. By evaluating macroevolutionary patterns within *Asplanchna*, we can better understand how a variety of novel traits emerge and evolve.

## Acknowledgements

We thank Karin Pfennig, Andrew Isdaner, and Sedona Kelly for feedback on experimental design.

