## Supplementary material for "Consequences of cannibalism: induced defense and kin discrimination in a rotifer": Table S1

**Supplemental Table**

**Table S1.** Average body lengths of *A. brightwellii* lineages

| Morph | Lineage | Neonates | Adults |
| --- | --- | --- | --- |
|  |  | Mean ± SD body length (µm) | |
| Beta | P | 495.0 ± 27.3 | 543.4 ± 78.1 |
|  | BRH | 454.6 ± 40.6 | 516.9 ± 75.1 |
|  | LT | - | 544.5 ± 80.6 |
|  | RS | - | 555.8 ± 68.8 |
| Alpha | 1P | 267.8 ± 30.9 | 383.8 ± 52.8 |
|  | 1LT | 283.3 ± 38.3 | 485.7 ± 65.9 |
|  | 2LT | - | 465.9 ± 93.0 |
|  | LP32 | - | 478.7 ± 83.6 |

BRH = Behind Ranch House, LT = Lonely Tank, P = Peccary Tank, RS=Rattlesnake Tank, W = Willow Tank
